# Development of a pipeline to evaluate YAP-TEAD inhibitor potential *-* direct TEAD inhibition represses cancerous growth in a *NF2*-mutant model of mesothelioma

**DOI:** 10.1101/2024.11.12.622964

**Authors:** Richard Cunningham, Siyang Jia, Krishna Purohit, Michaela Noskova Fairley, Yue Lin, Ning Sze Hui, Rebecca E. Graham, Adriano G. Rossi, Justyna Cholewa-Waclaw, Neil O. Carragher, Carsten Gram Hansen

## Abstract

As the core, tumorigenic downstream effectors of the Hippo signalling pathway, YAP/TAZ and the TEAD family of transcription factors represent attractive targets for drug discovery efforts within cancer research. This is particularly true within the context of pleural mesothelioma, in which there are many recent preclinical developments and clinical trials evaluating the efficacy of TEAD inhibitors. The range of inhibitors have shown great promise, but comparisons of their performances are so far limited. Here we develop a high content pipeline that enables a comparative analysis of currently developed YAP/TAZ-TEAD inhibitors. We take advantage of isogenic cellular models that enable us to examine inhibitor specificity. We identify genetic compensation of the Hippo pathway transcriptional module, with implications for therapeutic targeting, and implement Cell Painting to develop a detailed morphological profiling pipeline that enables further characterisation, quantification, and analysis of off-target effects. Our pipeline is scalable and allows us to establish specificity and comparative potency within cancer relevant assays in a clinically relevant cellular model.

## INTRODUCTION

Pleural mesothelioma is an asbestos induced cancer of the mesothelial lining of the lung and the deadliest cancer after diagnosis^1,2^. Patients have a bleak outlook as many are diagnosed late or with aggressive disease, limiting the efficacy of surgical intervention^3,4^. There are no curative therapies. To this end, therapeutics currently used can be considered palliative care and patients only survive on average for 12-18 months after diagnosis. This highlights a clear and urgent need for the development of targeted therapies and implementation of effective chemotherapeutics^5^. As a relatively uncommon cancer-type, pleural mesothelioma (PM) is defined by a lack of clear, high penetrance driver mutations^6–10^. Despite this, PM is distinct from most common cancers in that genomic perturbations within upstream components of the Hippo pathway are overrepresented within its mutational profile.

The Hippo pathway comprises an upstream kinase cascade module, which functions to regulate the activity of the co-transcriptional activators YAP and TAZ^11,12^. These, alongside the TEAD family of transcription factors^13,14^, represent the downstream transcriptional effectors of the Hippo pathway. Since the initial discovery of the signalling pathway as a core regulator of a variety of processes in development, differentiation, and regeneration^15,16^, YAP and TAZ are implicated as oncogenic drivers across a range of cancer-types^17–24^. Strikingly, PM is defined by a relatively high frequency of loss-of-function mutations within the upstream regulatory component of the Hippo pathway relative to other cancer types^6–10^. These mutations include frequent loss-of-function mutations within the upstream Hippo pathway kinase cascade, including most frequently in *NF2*, with more occasional loss-of-function mutations in *SAV1*, *LATS1* and *LATS2*^6–10^.

While designing therapeutics specifically against loss of tumour suppressors has been challenging historically, multiple inhibitors of the TEAD family of transcription factors have recently been developed with the hopes to position for clinical use. The realisation of the development of a range of varied direct inhibitors is an exciting advancement in cancer research, as hyperactive YAP/TAZ-TEAD activity is a widespread phenomenon across many cancers^25^. The majority of the inhibitors developed comprise small molecule disruptors of TEAD auto-palmitoylation, a post translational modification required for the interaction between these transcription factors and YAP/TAZ^26,27^. Following initial preclinical development, some of these promising inhibitors have now progressed to clinical trials, to be positioned as YAP/TAZ-TEAD inhibitors in PM patients. Due to the genomic evidence and frequent dysregulation of the Hippo pathway within PM, combined with both the ineffectiveness of surgery^28^ and general dearth of effective therapeutic interventions^10,29^, this cancer-type is the primary focus in the development of these therapeutics.

Beyond these selective inhibitors that are currently undergoing testing, there are a variety of indirect inhibitors of YAP/TAZ-TEAD. These comprise a heterogenous range of chemicals, including dasatinib, a Src kinase inhibitor commonly used as an anti-cancer agent^30,31^, verteporfin, a photosensitiser historically used as a chemical inhibitor of YAP/TAZ activity^32,33^, as well as statins^34^, a family of mevalonate pathway inhibitors reported to inhibit YAP/TAZ activity via the disruption of geranylgeranylation^19^. We focus on these different therapeutic classes and prioritize a range of inhibitors with a focus on those that have well-documented, potent anti-YAP/TAZ activity. By focusing on a range of both selective and classical, less selective inhibitors, we compare these side-by-side, taking advantage of two separate and complimentary isogenic cellular models to facilitate a direct comparison in the molecular impact of therapeutic families within a clean genetic background.

## RESULTS

### Identifying high potency inhibitors of YAP/TAZ-TEAD

With the current focus on YAP-TEAD inhibition in the development of chemotherapeutics for use, especially for the treatment of PM^29,35^, three recently developed TEAD inhibitors were selected for testing within cell-lines. This includes VT-107^36^ and K-975^37,38^, both pan-TEAD inhibitors, and IK-930, a TEAD1-specific inhibitor^39^ that has undergone clinical trials^40^ for the treatment of PM and other tumours associated with perturbations within the Hippo pathway (NCT05228015). A multitude of additional inhibitors of YAP-TAZ/TEAD have in the past been identified through screening approaches and used primarily as tool compounds to probe pre-clinical impacts of Hippo pathway perturbations, as well as undergoing further preclinical development and optimization^29,41^. Despite widespread use, most of these have less well-defined mechanisms of action (MoA) and might have YAP-TAZ/TEAD inhibition as a secondary and potential off-target effect^33,42^. To compare the inhibitory potential of recently developed, on-target TEAD inhibitors to these more classic YAP inhibitors, we included verteporfin, lovastatin, and dasatinib as comparators. Dasatinib is prioritized, as its primary mechanism-of-action as an anti-cancer therapeutic is via its function as a Src and tyrosine kinase inhibitor^30,31^, with Dasatinib’s downstream cellular impact on YAP/TAZ-TEAD appearing likely through the Src-YAP signalling axis^43,44^.

Initial testing was focused on the ability of compounds to inhibit cellular TEAD activity. This was analysed via the use of a luciferase reporter regulated by multiple TEAD binding sites^45^. Quantification of TEAD activity upon 24-hour treatment at 1 µM confirmed the ability of almost all tested compounds to inhibit TEAD activity (figure 1A), apart from lovastatin. Interestingly, dasatinib exhibits the most profound inhibition of TEAD activity. In order to validate this effect, we quantified the expression of YAP/TAZ signature genes, as defined by TCGA^46^, on treatment with a selection of these inhibitors. This reveals a modest, though significant, decrease in signature expression (figure 1B) under the same treatment conditions. Notably, the observed decrease in signature gene expression is TEAD-dependent, with no significant effect evident on treatment in TEAD KO cells^14^ (supplementary figure 1A). This suggests that not only TEAD inhibitors, but also off-target inhibitors such as verteporfin, modulate the expression of YAP/TAZ target genes in a TEAD-dependent manner. Given the difficulties associated in directly targeting YAP/TAZ directly due to their intrinsic disorder^29^, these findings reinforce the likelihood that YAP/TAZ may be primarily targetable via their interaction with TEADs^29^.

**Figure 1.**
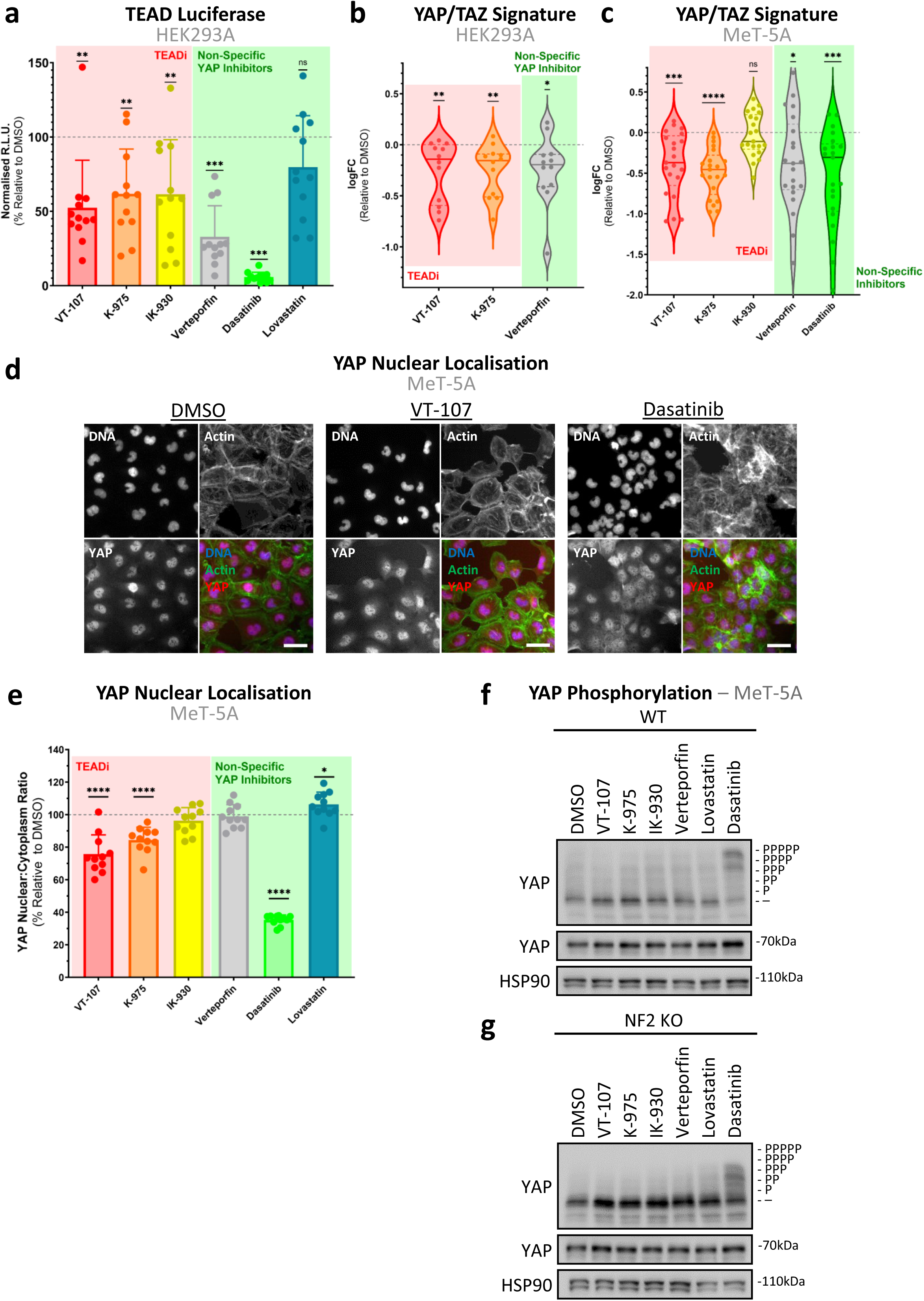
Direct inhibitors of TEAD and indirect inhibitors of YAP differentially effect YAP-TEAD activity *in vitro*. **a,** Bar-plot shows TEAD luciferase-based activity in HEK293A cells, treated at 1 µM with various selective TEADi (red), or non-specific YAP inhibitors (green) for 24 hours. Points represent individual technical replicates (n=4). All inhibitors, besides lovastatin, decrease TEAD based luciferase activity. **b,** Violin plot shows the expression of YAP/TAZ signature genes, as determined via RT-qPCR, in HEK293A cells after 24-hour treatment of select high potency inhibitors determined in (**a**) at 1 µM (n=3). Dots represent the log-transformed fold-changes of individual YAP/TAZ target genes, with a clear downregulation across most quantified signature genes. **c,** Violin plot, as in (**b**), shows downregulation of YAP/TAZ signature genes in MeT-5A cells on treatment with YAP/TAZ-TEAD inhibitors (n=3). **d,** Representative confocal images of YAP nuclear localisation in MeT-5A cells (DMSO, vehicle control, left), or with direct TEAD (VT-107, middle) or non-specific YAP (dasatinib, right) inhibitors at 370 nM for 24 hours. Increased cytoplasmic YAP is seen as compared to vehicle control (DMSO). Scale bars = 50 µm. **e**, Bar-plot shows quantification of images as depicted in (**d**), with data presented as % nuclear:cytoplasmic YAP ratios adjusted to vehicle controls. Each dot represents the median values of individual wells (n=3). **f,** Phos-Tag based western blot shows YAP phosphorylation in WT MeT-5A cells in response to 370 nM 24-hour treatment. YAP phosphorylation is induced solely on treatment with dasatinib. **g,** Phos-Tag based western blot conducted as in (**f**) on cell lysates from NF2 KO MeT-5A cells shows a decrease in sensitivity to dasatinib, in terms of YAP phosphorylation relative to WT cells. *P* values in (**a**)-(**c**) were determined by one-sample Wilcoxon signed rank test and (**e**) using Mann-Whitney U test, adjusted for multiple comparisons. n.s. = Not significant, *P < 0.05, **P < 0.01, ***P < 0.001, and ****P < 0.0001 relative to WT.

In order to probe the efficacy of candidate compounds within the context of mesothelioma, evaluation was then conducted in a preclinical model of PM driver mutations, with CRISPR-Cas9 mediated knockout^47^ (KO) of tumour suppressor NF2 in non-malignant, MeT-5A mesothelial cells. This unique isogenic cellular model represents the disease well^6^ and is an ideal platform for interrogating YAP/TAZ-TEAD dynamics and responses to therapeutic inhibition. We have previously shown that the TCGA-defined YAP/TAZ signature gene-set^46^ is upregulated on NF2 loss in the mesothelial context under cancer-related stresses^6^. Initial testing focused on the effect of treatment on YAP/TAZ-TEAD mediated transcription via quantifying the expression of these canonical downstream target genes. Computing the expression of this signature reveals that almost all tested inhibitors were sufficient to significantly downregulate the transcription of the full gene-set (figure 1C) after 24-hour treatment at 1 µM, reinforcing their abilities to suppress YAP/TAZ-TEAD activity. Interestingly, IK-930, a TEAD1-specific inhibitor, did not decrease signature expression under these conditions (figure 1C).

YAP/TAZ-TEAD activity is contingent on nuclear retention of YAP/TAZ, with the upstream kinase component of the Hippo pathway known to phosphorylate YAP/TAZ, leading to subsequent cytoplasmic sequestration and therefore inhibition of these co-transcriptional activators^48,49^. To assess the dynamics of YAP/TAZ alongside the observed reduction in YAP/TAZ-TEAD mediated transcription, we next quantified the inhibitory potential of candidate compounds across a range of concentrations in the context of YAP nuclear localisation. YAP was selected as the sole readout of activity here, with cellular TAZ dynamics omitted from study, due to YAP and not TAZ being the prime determinant of tumorigenicity within the MeT-5A cellular model^6^. Additionally, as inhibitors are presumed to be cytostatic/cytotoxic and YAP nuclear localisation is reduced under conditions of high cell density^29,50,51^, cells were filtered to those approximating 50% cell-cell contact to limit effects from contact inhibition at either extreme^6,51^. Although lower potency analogues of VT-107 and K-975 apparently do not impact nuclear YAP retention in mesothelioma cell-line models^36^ and the osteosarcoma U2OS cell-line^52^ respectively, we identified that the majority of selective TEAD inhibitors tested did modulate YAP localisation 24 hours post-treatment. This effect is observed by increased cytoplasmic/nuclear YAP ratio (figure 1D), indicating a reduction in the level of transcriptionally active YAP, with the maximal response generally observed at the lowest concentration point included (370 nM; supplementary figure 1B). Reduction in YAP nuclear localisation broadly matches decreased YAP/TAZ signature gene expression, with a clear effect observed when cells were treated with pan-TEAD inhibitors. However, in contrast to its effect on YAP-TEAD gene expression, there was no significant decrease in nuclear YAP with verteporfin treatment, with dasatinib being the prime non-selective inhibitor that induced cytoplasmic retention of YAP in MeT-5A cells (figure 1E).

With the inhibitory potential of these compounds established within MeT-5A cells, we next sought to assess this effect within the context of loss of the tumour suppressor *NF2*. Notably, loss of NF2 had little impact on sensitivity to YAP/TAZ-TEAD inhibition in terms of expression of YAP/TAZ signature genes, with just a slight decrease in sensitivity to K-975 observed (supplementary figure 1C), while NF2 KO cells exhibited slightly enhanced sensitivity to selective TEAD inhibition on YAP nuclear localisation (supplementary figure 1D). Though this effect was significant, it was modest, while sensitivity to dasatinib was conversely decreased in NF2 KO cells.

YAP’s principal nuclear/cytoplasmic regulation primarily takes place via LATS1/2 mediated inhibitive phosphorylation on five serine residues^49,53^. In order to interrogate the manner with which this YAP inhibition is mediated, we analysed cellular lysates after 24-hour treatment at 370 nM and 1 µM, performing Phos-Tag western-blotting, a sensitive technique that allows for the identification of the phosphorylation state of YAP^49,53^. This shows that dasatinib is the sole tested inhibitor that induces YAP phosphorylation at low concentrations (figure 1F), while lovastatin treatment results in YAP phosphorylation only at higher concentrations (supplementary figure 1E). This YAP phosphorylation indicates that dasatinib’s potent inhibitory effect on YAP/TAZ-TEAD (figure 1F), as well as lovastatin’s more modest effect (supplementary figure 1E), may be mediated via the core kinase module of the Hippo pathway. To explore this, we conducted Phos-Tag based western-blots using lysates from similarly treated LATS1/2 double knockout (dKO) HEK293A cells^54–56^. Cells lacking LATS1/2 have, as expected, less phosphorylated YAP, indicating nuclear and therefore hyperactive YAP (supplementary figure 1F). These cells also show no YAP phosphorylation response to treatment with lovastatin or dasatinib, while cells with intact LATS1/2 exhibit TEAD-independent phosphorylation of YAP on treatment (supplementary figure 1F). Combined, the results highlight that YAP phosphorylation induced by lovastatin or dasatinib treatment is dependent on LATS1/2 kinases and therefore likely mediated via the Hippo pathway kinase cascade.

Interestingly, the increase in phosphorylated YAP observed in NF2 KO MeT-5A cells upon dasatinib treatment is diminished relative to WT MeT-5A cells (figures 1F-G). This blunted response is consistent with the decreased sensitivity of NF2 KO cells in YAP nuclear localisation to dasatinib (supplementary figure 1D). Considered together, these findings suggest that NF2-loss may result in a resistance to inhibition of YAP/TAZ-TEAD via activation of the Hippo pathway’s core kinase cascade, while leaving NF2-deficient cells similarly vulnerable to TEAD inhibition as compared to those with intact NF2.

### YAP/TAZ-TEAD inhibition disrupts the tumorigenic potential of *NF2*-deficient mesothelial cells

With the activity of the candidate compounds established, we next sought to determine the *in vitro* anticancer potential of candidate therapeutics in the context of *NF2*-deficient mesothelioma. Initially, this was assessed by quantifying YAP/TAZ-TEAD inhibition’s impact on cellular viability. Proliferation assays performed over the course of 72 hours (figure 2A) revealed that selective TEAD inhibitors generally had little effect on cell growth (figure 2B). Only VT-107 showed a consistent, though modest, reduction over the course of 72 hours treatment. This was in stark contrast to non-selective inhibitors of YAP, which caused a complete cessation of growth at concentrations >1 µM (figure 2B).

**Figure 2.**
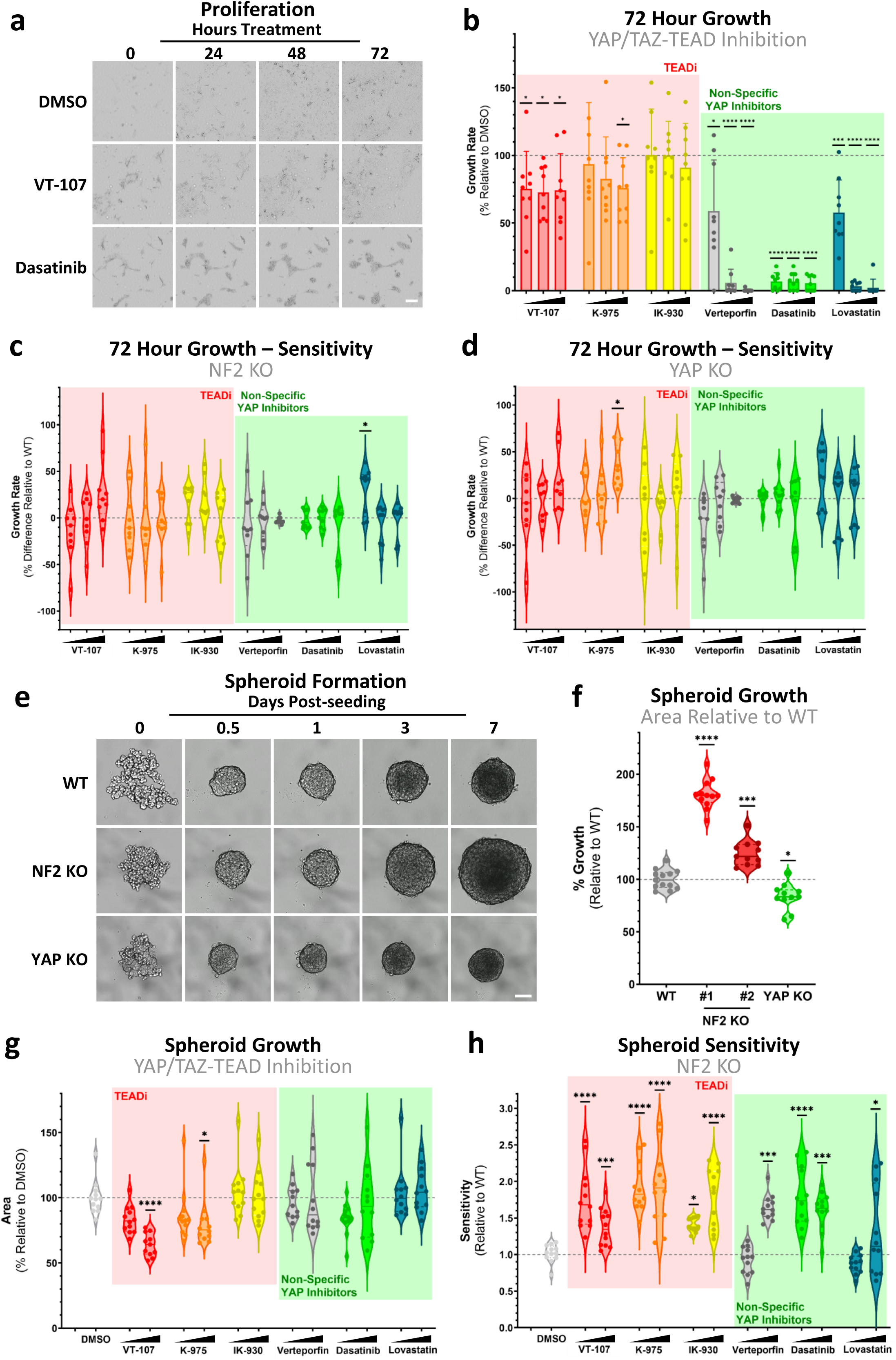
Specific TEAD-selective inhibition decreases NF2-deficient cancer-relevant phenotypes. **a,** Representative images of 2D proliferation in WT MeT-5A cells at 10 µM YAP-TEAD inhibition. Scale bar = 50 µm. **b,** Bar-plot of data from images as shown in (**a**) reveals inhibition of growth selectively in all non-specific YAP inhibitors (green) tested. Direct TEADi (red) has modest impact on cell growth in 2D compared to non-specific YAP inhibitors (green). Points represent the growth curve statistics of individual wells (n=3). **c,** Violin plot shows cell proliferative sensitivity of NF2 KO MeT-5A cells, with decreases in cell growth shown relative to those in WT cells (**b**). Little change in sensitivity is observed upon loss of NF2. **d,** Violin plot as in (**c**) showing sensitivity to YAP/TAZ-TEAD inhibition in YAP KO relative to WT MeT-5A cells, in terms of cell growth rates in 2D. **e,** Representative images of spheroid formation in WT (top), NF2 KO (middle) and YAP KO (lower). Scale bar = 50 µm. **f,** Quantification of spheroids imaged at day 7, as in (**e**), shown as violin plots. WT (grey), two NF2 KO clones (red), and YAP KO (green) MeT-5A cells. Genotypes form spheroids of varying sizes, with spheroids containing NF2 loss (larger) and YAP loss (smaller), respectively. #1 and #2 NF2 KO are independently generated clones. **g,** Violin plot shows sensitivity of WT MeT-5A spheroids to YAP-TEAD inhibition. Each dot represents size of an individual spheroid at day 7, relative to size at 24-hours to account for variable seeding (n=3). Spheroid size is modestly decreased upon high concentration TEAD inhibition. **h,** Violin plot, as in (**g**) shows sensitivity of NF2 KO spheroids to YAP-TEAD inhibition relative to WT. NF2 KO MeT-5A spheroids have enhanced sensitivity to most tested inhibitors. Inhibitors tested in (**b**)-(**d**) were used at 1.1, 3.3, and 10 µM and in (**g**)-(**h**) at 1 and 10 µM and are compared to vehicle control (DMSO). *P* values in (**b**)-(**h**) were determined by Mann-Whitney U test, adjusted for multiple comparisons. n.s. = Not significant, *P < 0.05, **P < 0.01, ***P < 0.001, and ****P < 0.0001 relative to WT.

On loss of NF2, sensitivities across all treatments remain almost equivalent to wildtype (WT) MeT-5A cells (figure 2C, supplementary figure 2A). This is somewhat unexpected, as recent findings have indicated that NF2-deficient mesothelioma cells are more sensitive to TEAD inhibitors^57^. Importantly, our studies are conducted on an isogenic background and are therefore not impacted by the complexities of comparing different heterogenous cell lines with varying epigenetic backgrounds and underlying genetic diversity beyond *NF2* status. Combined, these additional factors might modulate sensitivity to TEAD inhibition, obfuscating nuanced molecular dynamics. Interestingly, loss-of-viability induced by non-selective YAP inhibitors appears to be YAP-independent, with YAP KO MeT-5A cells exhibiting similar sensitivities as WT cells to all compounds tested (figure 2D). YAP-independence therefore might explain the initial observation that NF2 KO cells are equally sensitive to dasatinib in terms of proliferation (figure 2C, supplementary figure 2A), despite the relative insensitivity in terms of YAP activation (figures 1F-G).

Given the role of NF2 in regulating YAP/TAZ in response to mechanical stimuli, including contact inhibition, substrate stiffness, and the mechanotransduction mediated by the YAP/TAZ-TEAD axis, we next focused on the establishment of a functional assay capable of quantifying cellular response to these cancer-adjacent mechanical stresses. Previously, we identified NF2 as an upstream mechanoregulator in mesothelial cells, and that mesothelial NF2-loss is sufficient to drive YAP-mediated anchorage-independent growth^6^. As the soft agar assay is challenging to scale up for high-throughput assays, we sought to establish an analogue approach. We therefore implemented a spheroid formation assay to analyse the effects of the selected inhibitors in a scalable manner. When MeT-5A cells are cultured in an ultra-low attachment setting, they readily form spheroids, with spheroids exhibiting less nuclear YAP than cells cultured in 2D (supplementary figure 2B). Interestingly in spheroids, NF2-loss leads to relatively enhanced nuclear YAP signal (supplementary figure 2C), suggesting a resistance to the reduction in nuclear YAP observed in WT spheroids. To assess how these differences in YAP activity may drive distinct spheroid phenotypes, we compared the growth of spheroids with intact NF2 and YAP to those with either KO of NF2 or YAP (figure 2E). In addition to a relative increase in nuclear YAP, there is a concurrent increase in spheroid size in MeT-5A spheroids on loss of NF2 (figure 2F). In contrast, YAP-deficient spheroids exhibit a clear decrease in size (figure 2F). This increased growth observed in NF2-deficient spheroids recapitulates key aspects of the 3D tumour environment and therefore reflects tumorigenic potential^58^, possibly mediated via enhanced YAP activity.

Next, having established a quantifiable cancer-relevant phenotype robustly driven by NF2 loss, we tested spheroid growth in the presence of YAP/TAZ-TEAD inhibitors to quantify the impact of inhibition on the tumorigenic capacity of spheroids. WT MeT-5A cells exhibit partial sensitivity selectively to pan-TEAD inhibitors (figure 2G), with a significant reduction in spheroid growth observed on treatment with VT-107 and K-975 at high concentrations (10 µM). Interestingly, NF2 KO MeT-5A cells were more sensitive to nearly all treatments (figure 2H, supplementary figure 2D). Beyond sensitivity, our data highlight that NF2 KO cells exhibit a marked decrease in spheroid area, with an enhanced effect observed with on-target TEAD inhibition as compared to indirect YAP/TAZ-TEAD inhibitors (supplementary figure 2E). This inhibitory effect is sufficient to revert NF2-deficient spheroids to sizes equivalent to YAP KO spheroids.

These findings collectively point to the spheroid formation assay as effective to evaluate and probe YAP/TAZ-TEAD driven tumorigenic capacity.

### Determining the role of YAP and TEAD isoforms on YAP/TAZ-TEAD inhibitor efficacy

Given NF2-mutant *in vitro* anchorage-independent growth is dependent on YAP^6,51,59^, we next sought to establish the extent to which YAP-TEAD influence the observed NF2-deficient phenotype. To confirm YAP’s role in driving NF2 KO spheroid formation, YAP was overexpressed in WT MeT-5A cells. We exogenously expressed YAP-5SA, which contains point mutations across five LATS1/2 target residues, rendering YAP non-phosphorylatable by LATS, and therefore hyperactive^51^. Expression of YAP-5SA in MeT-5A cells (figure 3A) reveals that hyperactive YAP drives spheroid growth, closely phenocopying NF2 KO spheroids (figure 3B). To resolve whether this is driven by TEAD activity targetable via therapeutic inhibition, YAP-5SA spheroids were treated with TEAD-selective inhibitors. Spheroids with hyperactive YAP phenocopy the inhibition observed upon loss of NF2 KO, with enhanced sensitivity to TEAD inhibition (figure 2H, figure 3C).

**Figure 3.**
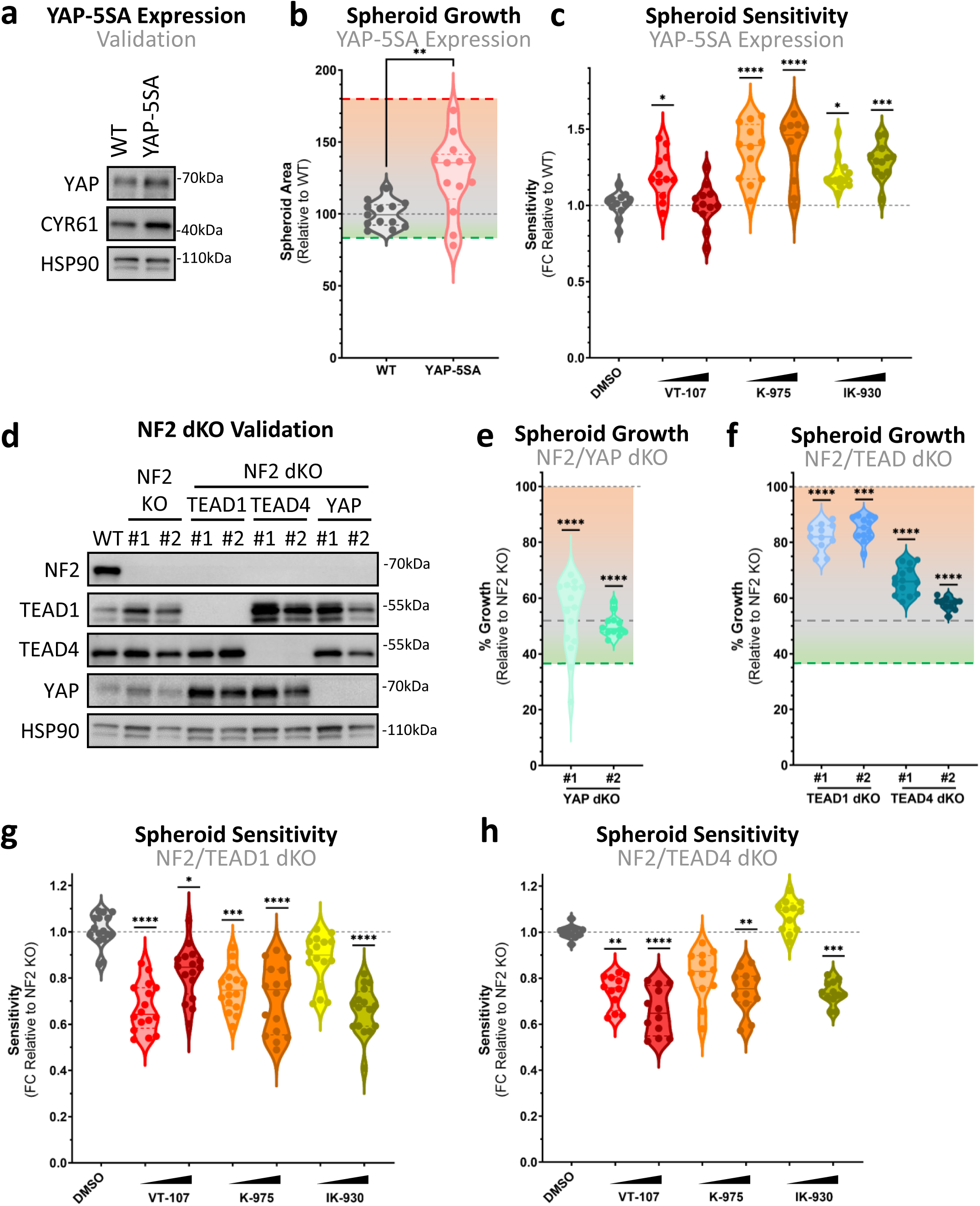
YAP and TEAD differentially orchestrate NF2-deficient spheroid sensitivity. **a,** Western blots from MeT-5A cells stably expressing YAP-5SA, a hyperactive form of YAP. An increase in protein levels of YAP and CYR61, a downstream transcriptional target of YAP-TEAD, confirm induction of YAP-5SA expression. **b,** Violin plot shows the relative spheroid growth of YAP-5SA expressing MeT-5A spheroids, compared to WT control. Gradients show % growth relative to parental NF2 single KO (red), WT (grey), and YAP KO (green). **c,** Violin plot shows sensitivity of MeT-5A spheroids expressing YAP-5SA to TEAD inhibition, relative to WT control spheroids. **d,** Western blots from gene edited combinatorial KO clones, show KO of YAP and TEAD isoforms alongside NF2 in MeT-5A cells. **e-f,** Quantification of relative spheroid growth shown in violin plot comparing impact of YAP (**e**) or TEAD (**f**) KO in NF2-deficient MeT-5A spheroids. Gradients as in (**b**) show % growth relative to single KO genotypes. **g-h,** Violin plots show sensitivity of NF2/TEAD1 dKO (**g**) and NF2/TEAD4 dKO (**h**) MeT-5A spheroids to TEAD inhibition. Loss of either TEAD isoforms results in a decrease in sensitivity to TEAD inhibition. Spheroids in (**c**), (**g**), and (**h**) were treated for 24 hours with 1 and 10 µM of selective TEAD inhibitors. Each dot represents an individual spheroid across 3 biological replicates. *P* values in (**b**), (**c**), (**e**)-(**h**) were determined by Dunnet’s multiple comparison test. n.s. = Not significant, *P < 0.05, **P < 0.01, ***P < 0.001 and ****P < 0.0001.

We next assessed spheroid growth in NF2/YAP double knockout (dKO) cells (figure 3D). When YAP is concurrently lost in NF2 KO cells, a reversal of the NF2 KO phenotype is observed, with NF2/YAP dKO spheroids exhibiting a dramatic decrease in spheroid growth relative to NF2 KO (figure 3E). As such, NF2/YAP dKO spheroid sizes were restored roughly to those observed in YAP KO spheroids. Taken together, these observations further reinforce the likelihood that YAP is the primary effector of NF2 KO spheroid growth and, given the enhanced sensitivity to TEAD inhibition observed in the presence of hyperactive YAP, may be targetable via inhibiting YAP-TEAD transcription.

The TEAD family consists of four proteins in vertebrae, in humans termed TEAD1-4^13^. Recent early-stage clinical evaluation has highlighted that pan-TEAD and TEAD isoform specificity might be a defining clinical feature in toxicity and efficacy^60,61^. The implications of isoform specific or pan-TEAD targeting in cancer is therefore of importance with pan-TEAD likely being more effective, but in general higher risk of toxicity. Assessing TEAD isoform functional importance and drug specificity is therefore critical during drug development to guide optimal clinical responses. These implications prompted us to generate NF2/TEAD isoform specific dKO cells for TEAD isoforms with antibodies available and where TEAD isoforms are robustly expressed and detected in MeT-5A cells. TEAD1 and TEAD4 were specifically recognised and targeted (figure 3D), with both TEAD1 and TEAD4 implicated in the progression and development of various cancers^62–64^. Notably, TEAD4 loss leads to an increase in protein levels of TEAD1, while loss of either TEAD1 or TEAD4 increases YAP levels (figure 3D). This finding has ramifications for mesothelioma, and potential general cancer therapy, given the possibility of compensatory upregulation to counter the inhibition of single components within the pathway. Loss of TEAD1 or TEAD4, in contrast to YAP loss, leads to a modest decrease in spheroid area (figure 3F), reverting spheroid size closer to that of WT spheroids. This effect is particularly pronounced in TEAD4 KO cells. When compared to NF2/YAP dKO, these results suggest complete and partial dependence on YAP and individual TEAD isoforms respectively.

To assess which of the four TEAD isoforms, if any, coordinate the hypersensitivity of NF2-deficient spheroids to TEAD inhibition, we treated NF2/TEAD1 and NF2/TEAD4 dKO MeT-5A spheroids and compared response to parental genotypes. If the observed response to TEAD inhibition is mediated via YAP-TEAD, a limited additive effect of spheroid reduction on combination of TEAD inhibition and KO of critical TEAD isoforms would be evident. Resulting dKO sensitivities indicate that, although TEAD4 may be more critical for the establishment of the observed NF2-deficient phenotype, with a more complete reversal on concurrent loss of TEAD4 as compared to TEAD1 (figure 3F), the loss of either isoform is sufficient to offset the increased sensitivity of NF2 KO spheroids to selective TEAD inhibition (figures 3G-H, supplementary figures 3A-B). More specifically, loss of TEAD1 leads to a blunting of NF2 KO spheroid sensitivity to selective TEAD inhibition, with all compounds exhibiting reduced sensitivity at one or both concentrations tested (figure 3G, supplementary figures 3A). Similar results are seen when TEAD4 is lost (figure 3H, supplementary figures 3B), which is particularly striking given IK-930 is purported to be a TEAD1-selective inhibitor. The decrease in sensitivity observed in NF2/TEAD4 dKO spheroids suggests that TEAD4 may coordinate, at least in part, some response to IK-930 (figure 3H).

In summary, these findings validate the role of YAP and TEADs in coordinating the tumorigenic capacity of NF2 loss *in vitro*, with YAP in particular indispensable to the enhanced spheroid forming capability of NF2-deficient cells. Though there is likely some redundancy among the TEAD isoforms, the enhanced response to TEAD inhibition in NF2-deficient spheroids is blunted when TEAD isoforms are concurrently eliminated.

### Morphometric profiling predicts selectivity of YAP/TAZ-TEAD inhibitors

Drugs with off-target effects are a major issue for drug development, directly impacting multiple levels of clinical transition, with common issues including toxicity, specificity, and efficacy affecting progress^65–68^. Treatment with inhibitors that exhibit broad off-target effects often induce morphological changes beyond their effect on proliferation, stemness, and therapeutic resistance. This is exemplified in the disruption of actin organisation in MeT-5A cells on treatment with dasatinib (figure 1D), consistent with previous observations across multiple cell-lines^30,69^. Additionally, a highly disruptive morphology is observed in spheroids on treatment with high concentrations of non-selective inhibitors (supplementary figure 3C), pointing to a recurrent phenotype associated with off-target effects of YAP/TAZ-TEAD inhibition.

As shifts in cellular morphology are predictive of broad pharmacological effects^70^ and can be leveraged to infer detailed molecular effects of treatment, we next sought to quantify the morphological disruption on YAP/TAZ-TEAD inhibition in our cellular model system. To initially confirm the phenotype observed in MeT-5A spheroids (supplementary figure 3C), we performed a quantitative analysis of the morphological shift of spheroids on treatment with YAP/TAZ-TEAD inhibitors. Brightfield images were segmented, and morphometric characteristics extracted and processed according to Joint Undertaking in Morphological Profiling (JUMP) pipelines^71^. This revealed a distinct morphological clustering within non-specific YAP inhibitors, while selective TEAD inhibition results in a phenotype that readily clustered with vehicle control cells (supplementary figure 3D).

Cell Painting is a newly developed approach to morphological profiling, which has been incorporated into high-throughput imaging assay pipelines to impute mechanisms-of-action^72^, bioactivity^73^, and toxicity^74,75^ in recent drug discovery efforts. In order to quantify the morphological perturbations induced on inhibition of YAP/TAZ-TEAD in greater resolution, we undertook Cell Painting to generate high-dimensionality datasets. Cell Painting involves the staining of cells with six dyes (supplementary figure 3E) and was performed in accordance with protocols established by the Joint Undertaking in Morphological Profiling Cell Painting (JUMP-CP) consortium^76^. Principal component analysis (PCA) of high-dimensional morphometric data reveals a morphological shift on treatment with non-selective inhibitors of YAP (figure 4A). To quantify this shift, the correlation distance across all morphological features was computed between treated wells. The resulting correlation distance matrix shows a clustering of non-selective and high concentration inhibitors, which share a high degree of morphological similarity (figure 4B). Interestingly, there was little clustering in vehicle control and low concentration TEAD-selective inhibitor wells. This suggests that morphological uniformity may be a metric of non-specific inhibition of YAP/TAZ-TEAD. Quantification of uniformity, as defined by the mean correlation distance between like-treated cells, reveals significantly enhanced morphological uniformity on high concentration, as well as dasatinib and lovastatin, treatment (figure 4C). To assess whether this observed uniformity is YAP-dependent or - independent, we quantified morphological uniformity on treatment in YAP KO MeT-5A cells (figure 4D). This reveals similar trends, with both dasatinib and lovastatin inducing a shift to more morphologically similar cells, thus indicating YAP-independence. Strikingly, high concentration K-975 and verteporfin treatment induced no such effect in YAP-deficient cells in contrast to WT, which may suggest a degree of morphological disruption mediated via YAP-TEAD inhibition.

**Figure 4.**
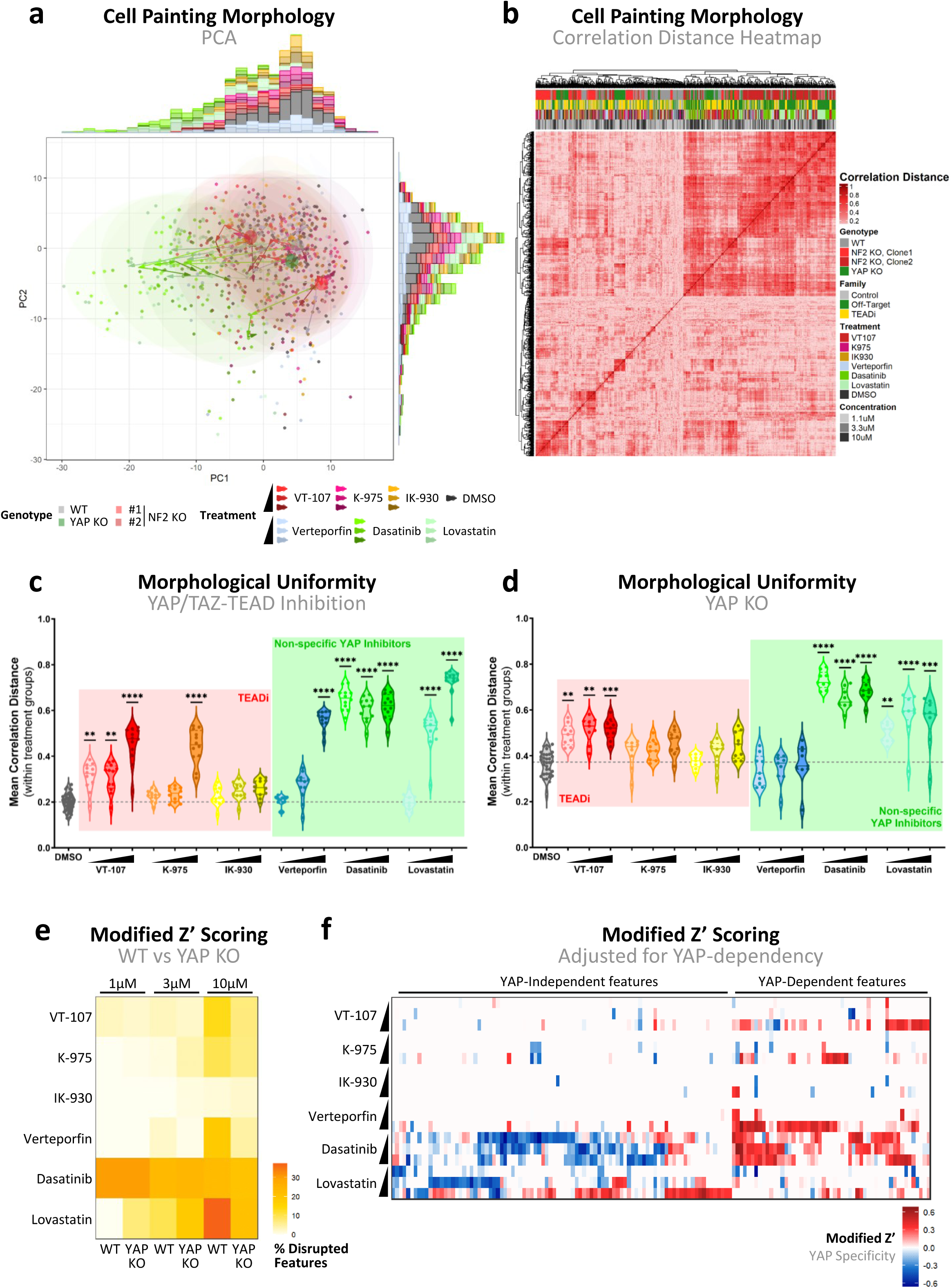
Morphological disruption predicts specificity of YAP/TAZ-TEAD inhibitors. **a,** PCA plot shows morphological profiles of MeT-5A cells cultured in 2D, with features collapsed via dimensionality reduction to allow visualisation. Each point represents a single well, with highlighted circles representing the centre of DMSO treated cells coloured according to genotype. Ellipses show the spread of treatments irrespective of genotype and arrows show trajectories across increasing concentrations of treatment for each genotype. **b,** Heatmap shows correlation distance of computed features between wells. Clustering of cells, each representing a single well, was performed via complete linkage, with annotations (top) showing the genotype, inhibitor family, name of treatment, and treatment concentration for each well. **c,** Violin plot portraying morphological uniformity of WT MeT-5A cells on YAP/TAZ-TEAD inhibition. Each point represents a single well, with uniformity calculated as the mean correlation distance of a single well to all like-treated wells. **d,** Violin plot as in (**c**), showing morphological uniformity on YAP/TAZ-TEAD inhibition in YAP KO MeT-5A cells. **e,** Heatmap shows the percentage of disrupted features, as defined by modified Z’ scores > 0, in WT vs YAP KO MeT-5A cells in response to various YAP/TAZ-TEAD inhibitors. As treatment concentration increases, a greater percentage of features are disrupted across most inhibitors. **f,** Heatmap shows modified Z’ scores of features with known discriminatory potential, defined as those with modified Z’ scores > 0, adjusted for YAP-dependency. Positive values indicate feature disruption selectively in WT, and not in YAP KO MeT-5A cells, indicating dependence on YAP, while negative values show features disrupted to the same or greater degree in YAP KO cells. Cells in (**a**), (**c**), (**d**), and (**f**) were treated for 24 hours with 1.1, 3.3, and 10 µM of compounds. *P* values in (**c**) and (**d**) were determined by Dunnet’s multiple comparison test. n.s. = Not significant, *P < 0.05, **P < 0.01, ***P < 0.001 and ****P < 0.0001.

With the broad impact of treatment on cell morphology established, we next performed in-depth analysis of specific features disrupted on inhibition of YAP/TAZ-TEAD. For this, we implemented a modified robust Z’ scoring, a parameter typically used in quality control of drug screening to extract features for each treatment that resolve distinctly from untreated controls. The Z’ scoring reveals a disruption of a high percentage of features on treatment with non-specific YAP and high concentrations of TEAD-specific inhibitors (figure 4E), consistent with broad profile analysis. Interestingly, these features are generally equally disrupted in YAP KO cells, particularly in dasatinib treated cells (figure 4E), suggesting that this morphological disruption is YAP-independent. To quantify this YAP specificity in detail, we adjusted modified Z’ statistics in WT to YAP KO cells. This revealed a range of conserved features that were consistently disrupted in a YAP independent manner specifically on treatment with the non-specific YAP inhibitors dasatinib and lovastatin (Figure 3F). Conversely, a smaller proportion of features were YAP dependent, which were disrupted on treatment with both selective TEAD inhibitors, with minimal overlap between the various TEAD inhibitors, and non-specific YAP inhibitors. Intriguingly, this suggests that different sets of features may be used to quantify YAP-dependent and -independent morphological changes, with verteporfin and both selective TEAD inhibitors showing less off-target YAP effects than dasatinib and lovastatin.

## DISCUSSION

Given the critical role the Hippo pathway and, more specifically, its downstream effectors YAP/TAZ-TEAD, play in the progression of mesothelioma and other cancers, there is a clear need to develop pipelines capable of assessing the efficacy and specificity of newly developed inhibitors to streamline positioning to clinical use. While *in vitro* studies assessing the efficacy of TEAD inhibition in NF2-deficient mesothelioma cells have been described^57^, these are limited by the heterogenous genomic and transcriptomic landscapes associated with cell-lines of mixed origins, potentially obfuscating critical, nuanced molecular dynamics of inhibiting of the transcriptional machinery of the Hippo pathway. Here, we present a pipeline within an isogenic model of loss of upstream and downstream components within the Hippo pathway, allowing the detailed interrogation of the molecular interplay of the individual constituents that comprise this tumorigenic pathway. We additionally describe functional assays for quantifying the YAP/TAZ-TEAD selective inhibition of tumorigenic capacity, which have shed light into the specifics of efficacy and selectivity of the core compound library used to target the major players within the pathway. Prior multiomic and genetic analysis have highlighted the *in vivo* complexities of pleural mesothelioma^6–9,77,78^. The unfortunate continued lack of curative therapies for this deadly disease is likely due to late-stage diagnosis and the sparsity of complimentary preclinical models representing the disease required for the predictive evaluation of drug targets before clinical evaluation in patients. Notably, our observations predict that targeting the transcriptional complex directly via inhibition of TEAD, including via its auto-palmitoylation site, is more specific, and thereby likely more effective, than targeting the upstream kinase module. Our observations also correlate well with clinical data, including the lack of activity and overall unwelcome association with pulmonary toxicities with dasatanib treatment in unselected pleural mesothelioma patients^79^. Our platform combines an extensive, *in vitro* based isogenic cellular model allowing for specificity evaluation, cancer-relevant assays where tumour suppressors are functionally active, high content imaging, and Cell Painting. Collectively, these allow for direct evaluation of YAP/TAZ-TEAD(1-4) targeting drugs and is feasible to implement alongside synergistic combinatorial drug evaluations. We are hopeful, that our approach provides a milestone within the development of mesothelioma treatments, as we appear to potentially evaluate potency of inhibition^80^ and clinical efficiency. The development of the platform thereby might inform future clinical trials and overall complement ongoing efforts to develop advanced cellular models^81,82^ and ultimately curative therapies.

Results point to K-975 as exemplar inhibitor of TEAD auto-palmitoylation, with VT-107 exhibiting equivalent potency alongside off-target effects at higher concentrations. Conversely, while IK-930 treatment is not associated with off-target effects and therefore a favourable safety profile, its potency is markedly less relative to alternative TEAD inhibitors. Despite previously reported selectivity for TEAD1^83^, the observation that loss of TEAD4 impacts the sensitivity of spheroids to IK-930 to the same extent of TEAD1 indicates some degree of inhibition of additional TEAD isoforms. Our findings also highlight the apparent compensatory mechanisms in place when targeting individual components (figure 3D). This discovery may explain the promiscuity observed in IK-930, suggesting a compensatory upregulation of, and therefore potential dependence on, TEAD1 on loss of TEAD4. Interestingly our results echo preliminary results from clinical studies of IK-930, which indicate that, despite being well tolerated, IK-930 has not proved effective in patients enrolled in clinical trials^84^.

In comparison to selective TEAD inhibitors, upstream inhibitors of YAP/TAZ-TEAD are associated with sweeping off-target effects, with dasatinib inducing cellular changes well beyond inhibition of the YAP-TEAD axis alone. Surprisingly, given the extent of YAP-independent effects previously associated with treatment, verteporfin exhibits a high degree of YAP-specificity across a range of functional assays. The convergence between phenotypes in cells treated with verteporfin and selective inhibitors of TEAD reinforces previous findings that verteporfin can act to disrupt YAP-TEAD binding^32^, though YAP-independent cytotoxicity is validated within our model system. The observation that high potency inhibitors of TEAD auto-palmitoylation also increases cytoplasmic retention of YAP is novel, though consistent with reports that inhibitors of YAP-TEAD interactions concomitantly reduce levels of nuclear YAP in mesothelioma cell-lines^85^. This phenomenon has been described as likely a result of displacement of YAP from chromatin-resident TEAD, reinforced by the finding that K-975 treatment reduces formation of YAP nuclear condensates, indicative of decreased binding to super-enhancer regions^52^.

To conclude, this work provides a framework for quantifying the efficacy of YAP/TAZ-TEAD inhibition for the treatment of mesothelioma. Conventional phenotypic screening approaches, which typically gauge success based on the cytotoxic or cytostatic properties of therapeutics, are impeded by their limited scope and are biased to drugs with broad toxicity profiles^86^. As the Hippo pathway integrates a wide range of stimuli^11,12^, it might be particularly prone to these perturbations. The pipeline established here benefits from the incorporation of a confirmed YAP-TEAD specific phenotypic assay, while the identification of a feature signature of YAP-specificity allows for the assessment of potential off-target effects of putative treatments. We expect this pipeline may help expedite the discovery of therapeutics effective in managing YAP/TAZ-TEAD driven mesothelioma, mitigating some of the risks of moving into *in vivo* assessment of inhibitor safety and efficacy.

## AUTHOR CONTRIBUTIONS

RC: data curation, conceptualization, formal analysis, validation, investigation, visualization, methodology, and writing—original draft, review, and editing. SJ: data curation, formal analysis, validation, investigation, visualization, methodology, and writing—review and editing. KP: data curation, formal analysis, validation, investigation, visualization and methodology. JCW: methodology and resources. MNF: investigation. YL: validation and methodology. NSH: investigation. REG: methodology. AGR: supervision. NOC: resources, methodology, review and editing. CGH: conceptualization, resources, data curation, formal analysis, supervision, investigation, methodology, and writing—original draft, review, and editing.

## ACKNOWLEDGEMENTS

Ongoing research in the Hansen lab (CGH) was funded by Worldwide Cancer Research (19-0238) and CSO-LifeArc. This project was initiated by pump prime funding from ISSF3 and JHMRF. MN and KR are funded by MRC Precision Medicine DTP Studentships. Nancy by the Martin Lee DTP. SJ is funded by a scholarship from the Chinese Scholarship Council, and the Edinburgh Global from University of Edinburgh. We furthermore acknowledge team members for insightful comments and constructive feedback during this study. We acknowledge the technical support and guidance provided by the Institute for Regeneration and Repair (IRR) Flow Cytometry and Cell Sorting Facility staff. Single cell sorting necessary for clonal selection of expansion of KO cell populations was conducted with support from the QMRI Flow Cytometry and IRR Flow Cytometry and Cell Sorting Facility, University of Edinburgh.

## MATERIALS AND METHODS

### Luciferase assay

HEK293A cells were seeded in triplicate in 12-well plates before co-transfection with Gal4-TEAD, 5×UAS Luciferase reporter, and *Renilla* luciferase constructs, using GenJet transfection reagent (SL100488, SignaGen). Post-transfection, cells were lysed and luciferase quantification was performed using a Dual-Glo® Luciferase Assay kit (E2920, Promega) according to manufacturer’s specifications and measured on a Biotek Synergy HT plate reader, with luminescence adjusted to *Renilla* luciferase.

### Culture maintenance

MeT-5A mesothelial cells and CRISPR-Cas9 mediated KO genotypes were generated and cultured as described in Cunningham *et al*^6^. HEK293A gene edited cells are described in Hansen *et al*^87^, Rausch *et al*^88^, and Lin *et al*^14^ and cultured in high glucose DMEM (21969-035, Gibco) with 2 mM L-Glutamine (25030-024, Gibco) and 10% FBS (10500-064, Gibco). Cells were grown in the presence of 100 units/mL of penicillin and 100 µg/mL of streptomycin (15140-122, Gibco) and incubated at 37°C with 20% O_2_. Where specified, cells were treated with VT-107 (HY-134957, MedChemExpress), K-975 (HY-138565, MedChemExpress), IK-930, verteporfin (SML0534, Sigma Aldrich), lovastatin (S2061-SEL, Selleckchem), and dasatinib, gifted by Prof Neil Carragher.

### Gene knockout and ectopic expression of YAP-5SA

Briefly, CRISPR-Cas9 mediated KO was carried out in MeT-5A mesothelial cells as described in Cunningham *et al*^6^, with the additional use of the following guide RNA (gRNA): 5’-TGGCAGTGGCCGAGACGATC-3’ for TEAD1 and 5’-CTCAAGGATCTCTTCGAACG-3’ for TEAD4. Validation of KO via western blotting. Expression of YAP-5SA in MeT-5A cells was performed by lentiviral transduction via pQX system under hygromycin B (30-240-CR, Corning) selection. Lentivirus for transduction was produced in HEK293T cells, harvested 48 and 72 hours after transfection using GenJet transfection reagent (SL100488, SignaGen), and filtered using low binding 0.45 µm SFCA filters (Corning, 431220). Selection was carried out via hygromycin treatment 24 hours post-transduction to ensure time for the development of resistance.

### Western Blotting

Western blots were performed with cell lysates harvested and run using homecast gels, as described in Hansen *et al*^87^. PageRuler prestained Protein Ladder (26616, Thermo Scientific) was included in western blots as a scale for protein size. Separated proteins were subsequently transferred from gels to Immobilon-P PVDF membranes (IPVH00010, Millipore) and blocked in 5% milk in TBS-T, with subsequent primary and secondary antibody incubation, washing, and development carried out as described in Cunningham *et al*^6^. Phos-Tag western blots were conducted with Phos-Tag reagent (304-93521, Alpha Laboratories) and 10mM MnCl_2_ added to each polyacrylamide resolving gel. Primary antibodies used were as follows: YAP (ab52771, Abcam), CYR61 (14479, CST), NF2 (D1D8, CST), TEAD1 (610922, BD Biosciences), TEAD4 (ab58310, Abcam), with HSP90 (BD610418, BD Biosciences) used as a loading control for samples.

### RT-qPCR analysis

Cells were plated and harvested, with cDNA generated as described in Cunningham *et al*^6^. RT-qPCR assays were carried out using Brilliant III Ultra-Fast SYBR Green (600883, Agilent) and LightCycler 480 SYBR Green I (04707516001, Roche) RT-qPCR Master Mixes, used according to manufacturer’s directions. IDT primers were custom-designed using templates deposited on PrimerBank^89^. RT-qPCR was carried out on a QuantStudio 5 Real-Time PCR System, with resulting data analysed using R statistical software. Primer sequences were as follows: 5’- GCTCGTTGAGTGAACGGCT-3’ and 5’- CATGAGCTAGTACAACATGAGGG -3’ for *AMOTL2*; 5’- AGTAGAGGAACTGGTCACTGG-3’ and 5’-TGTTTCTCGCTTTTCCACTGTT-3’ for *ANKRD1*; 5’- CCAAGGTGAGCTTTCCCTCG-3’ and 5’-CCTACTAGACCATAGGTCGTCGT-3’ for *ARHGEF17*; 5’- TAGAACAGCCCTTCAGAAAGTGA-3’ and 5’-CGGGGTTGTCTCGACTTAAAAA-3’ for *ASAP1*; 5’- GTGGGCAACCCAGGGAATATC-3’ and 5’-GTACTGTCCCGTGTCGGAAAG-3’ for *AXL*; 5’- CCCTGTGACGAGTCCAAGTG-3’ and 5’-GGTTCCGTAAATCCCGAAGGT-3’ for *CRIM1*; 5’- GAGGCAGAAGTACGGGGTTG-3’ and 5’-CAGGAATCACGGTTTCATGCT 3’ for *DOCK5*; 5’- GGCGCTTCAGGCACTACAA-3’ and 5’-TTGATTGACGGGTTTGGGTTC-3’ for *F3*; 5’- GCTGGTGGACCTAGTACAATGG-3’ and 5’-CTTACGAGCCGGTCGAAGTTG-3’ for *FJX1*; 5’- AATGCCACTCGCCCTACAC-3’ and 5’-CGTTCTGGTGCAAGTAGCTCT-3’ for *FOXF2*; 5’- GAGAGCAGAAGACCGAAAGGA-3’ and 5’-CACAACACCACGTTATCGGG-3’ for *GADD45A*; 5’- AGAGCACAGATACCCAGAACT-3’ and 5’-GGTGATTCAGTGTGTCTTCCATT-3’ for *IGFBP3*; 5’- ACTTTTCCTGCCACGACTTATTC-3’ and 5’-GATGGCTGTTTTAACCCCTCA-3’ for *LATS2*; 5’- TAATTGGCACGGCGACTGTAG-3’ and 5’-GGAGATCAGCTTGTACGGCAG-3’ for *MYOF*; 5’- GCCTGGGAGCTTACGATTTTG-3’ and 5’-TAGTGCCCTGGTACTGGTCG-3’ for *NT5E*; 5’- CGCCCAAGCCCCTAATGAAG-3’ and 5’-TCCCTCCGTATGTGCATCAGA-3’ for *NUAK2*; 5’- ATGCCTTTTGGTCTGAAGCTC-3’ and 5’-CCCTGTGCTTTCCACCGAC-3’ for *PTPN14*; 5’- GGGGAACAGTTGAGTAAAACCA-3’ and 5’-ACAATTTTTCCATACGGTTGGCA-3’ for *RBMS3*; and 5’-CAGCACACTCGATATGGACCA-3’ and 5’-CCTCGGGCTCAGGATAGTCT-3’ for *TGFB2*. All gene expression values were normalized to *Hypoxanthine Phosphoribosyltransferase 1* (*HPRT1*) expression.

### YAP nuclear localisation high content image-based assays

Cells were seeded in 384-well µClear plates (781091, Grenier) at variable cell densities determined for each genotype to ensure cells equivalent confluency over time-points tested. WT MeT-5A cells were seeded at 1,000 cells per well, NF2 KO at 833 cells per well and YAP KO at 2,000 cells per well. Upon attachment, cells were treated with relevant inhibitor for 24 hours before fixing, permeabilising, and staining as described in Cunningham *et al*^6^. Fluorescent cells were imaged using the Opera Phenix Plus high-content imaging system (PerkinElmer) at 20x across 3 biological replicates, with 9 fields imaged per well and four wells per sample/condition to act as technical controls. A bespoke CellProfiler^90^ pipeline was implemented to segment cells and quantify relevant intensities and morphological features. Statistical analyses were conducted using R, with cell-level data trimmed to ensure detectable nuclear and cytoplasmic YAP. Cells were trimmed according to cell-contact, with those between 45-55% cell-cell contact retained (7.6% cells of 1,381,549 total, with a median of 89 cells/well included for downstream analysis) and variance accounted for post-collation by removing outliers, defined as wells possessing values greater than 2 median absolute deviations (MAD) from the median of biological replicates. Processed data were plotted with GraphPad Prism.

### Proliferation and spheroid formation assays

For quantification of proliferation, cells were seeded at 2,000, 1,500, or 2,500 cells per well for WT, NF2 KO and YAP KO MeT-5A cells respectively in 96-well µClear plates (655090, Grenier) and treated with relevant inhibitors when attached, 24-hours post-seeding. Cells were then imaged over the course of 72-hours and confluency calculated using the CELLCYTE X (CYTENA). Growth was quantified by normalising slope statistics from computed logistic growth curves to DMSO treated wells, with rates of growth < 0 adjusted to 0. For spheroid formation assays, cells were seeded at 500 cells per well in ultra-low attachment 96-well plates (CLS7007, Corning) and cultured for 7 days. For live imaging assays and quantification of spheroid size, cells were incubated and imaged throughout this time period using the CELLCYTE X. To limit the influence of technical artefacts associated with the high variance of this platform, time-points exhibiting spheroid areas greater than 2 MADs from the median area at that time-point were trimmed, with any well with >50% trimmed time-points defined as outliers and excluded from analysis. 7-day spheroid areas were normalised to 24-hour time-points within each well. For treated spheroids, treatment was initiated ∼1 hour post-seeding, with resulting spheroids imaged at day 7 using the EVOS FL Auto 2 Imaging System (Invitrogen) at 10x magnification. Quantification of spheroid size was then achieved using CellProfiler, with the CellPose^91^ plugin to enhance spheroid segmentation. Brightfield images acquired were then subjected to morphological analysis as described in Cell Painting and morphological analysis, with a pipeline modified for single-channel quantification.

### Cell Painting and morphological analysis

Cell Painting assays were conducted according to established protocols^76^, with cells seeded as with high-content imaging in 384-well µClear plates (781091, Grenier) and treated with appropriate inhibitor for 24 hours before fixing, staining and imaging. Image acquisition using the Opera Phenix Plus high-content imaging system (PerkinElmer) was performed with the same set-up as described for YAP nuclear localisation high content image-based assays. Morphometric datasets were then acquired from resulting images using CellProfiler^90^ pipelines established by JUMP-CP protocols^71,92^, with the implementation of a modified version of pipelines available at github.com/broadinstitute/imaging-platform-pipelines/tree/master/JUMP_production. The cytominer^93^ package was then used to trim low variance or highly correlated features (leading to retention of 417 of 3,446 input features; 12.11%) and normalise retained features to vehicle control (DMSO) treated WT MeT-5A cells. Normalised high-dimensional morphometric datasets were visualised using circlize^94^ package in R. Morphological uniformity was computed by calculating correlation distances between all preserved features in each like-treated well. Feature disruption was quantified via a modified robust Z’ scoring, designed to act as a simplified, relative measure of a feature to resolve treated cell morphology from vehicle control (DMSO). This was calculated by removing the scaling constant of 3 of the traditional Z’ score and using median/MAD as opposed to mean/standard deviation as measures across wells, given no assumption of data to conform to a normal distribution. To implement scoring, the following equation was used:

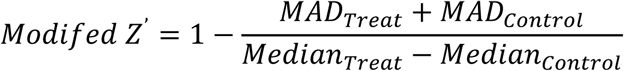

Modified Z’ scores were adjusted to quantify YAP dependency by subtracting YAP KO from WT scores.

---

**Supplementary Figure 1.**
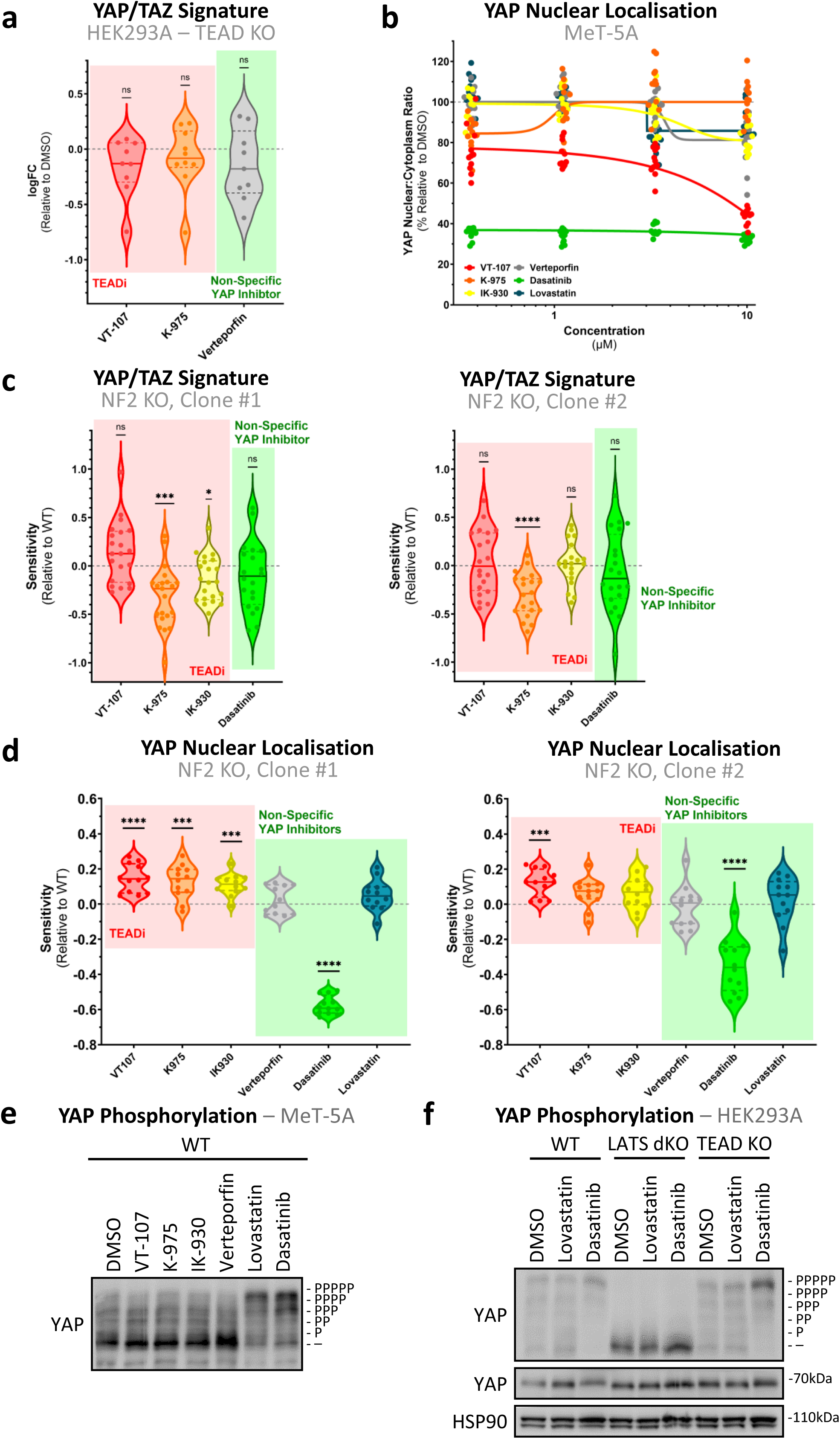
TEAD, but not NF2, mediates effects of YAP/TAZ-TEAD inhibitors. **a,** Violin plot as in figure 1b shows the expression of YAP/TAZ signature genes, as determined via RT-qPCR, in TEAD KO HEK293A cells, with expression depicted as log-transformed fold-change on treatment with select YAP/TAZ-TEAD inhibitors relative to DMSO. No significant dysregulation of signature genes is observed on treatment. **b,** Dose response quantification of decrease in nuclear YAP from images as in figure 1d. Dots represent the median nuclear:cytoplasmic YAP ratio across cells in individual wells over 3 biological replicates after 24-hour treatment. **c,** Violin plots comparing sensitivity of NF2 KO relative to WT MeT-5A cells to YAP/TAZ-TEAD inhibition, in the context of YAP/TAZ signature expression. Dots represent the individual genes comprising the signature, where sensitivity is the fold-change in up/down-regulation relative to WT response in two independent NF2 KO clones (left and right). **d,** Violin plots as in (**c**) show the sensitivity of NF2 KO MeT-5A cells to inhibition in the context of YAP nuclear localisation in two independent NF2 KO clones (left and right). **e,** Phos-tag based western blot, as in figure 1f, shows YAP phosphorylation in WT MeT-5A cells in response to 24-hour treatment with YAP/TAZ-TEAD inhibitors at 10 µM. **f,** Phos-tag based western blot (top) showing difference in response between WT (left), LATS1/2 dKO (mid), and TEADs KO (right) HEK293A cells to YAP/TAZ-TEAD inhibition, in terms of difference in YAP phosphorylation. The same lysates were analysed on a conventional Western blot and probed for YAP and HSP90 (bottom). Where not stated, cells were treated with 1 µM inhibitor for 24-hours. *P* values in (**a**) and (**c**) were determined by one-sample Wilcoxon signed rank test and (**d**) using Mann-Whitney U test, adjusted for multiple comparisons. n.s. = Not significant, *P < 0.05, **P < 0.01, ***P < 0.001, and ****P < 0.0001 relative to WT.

**Supplementary Figure 2.**
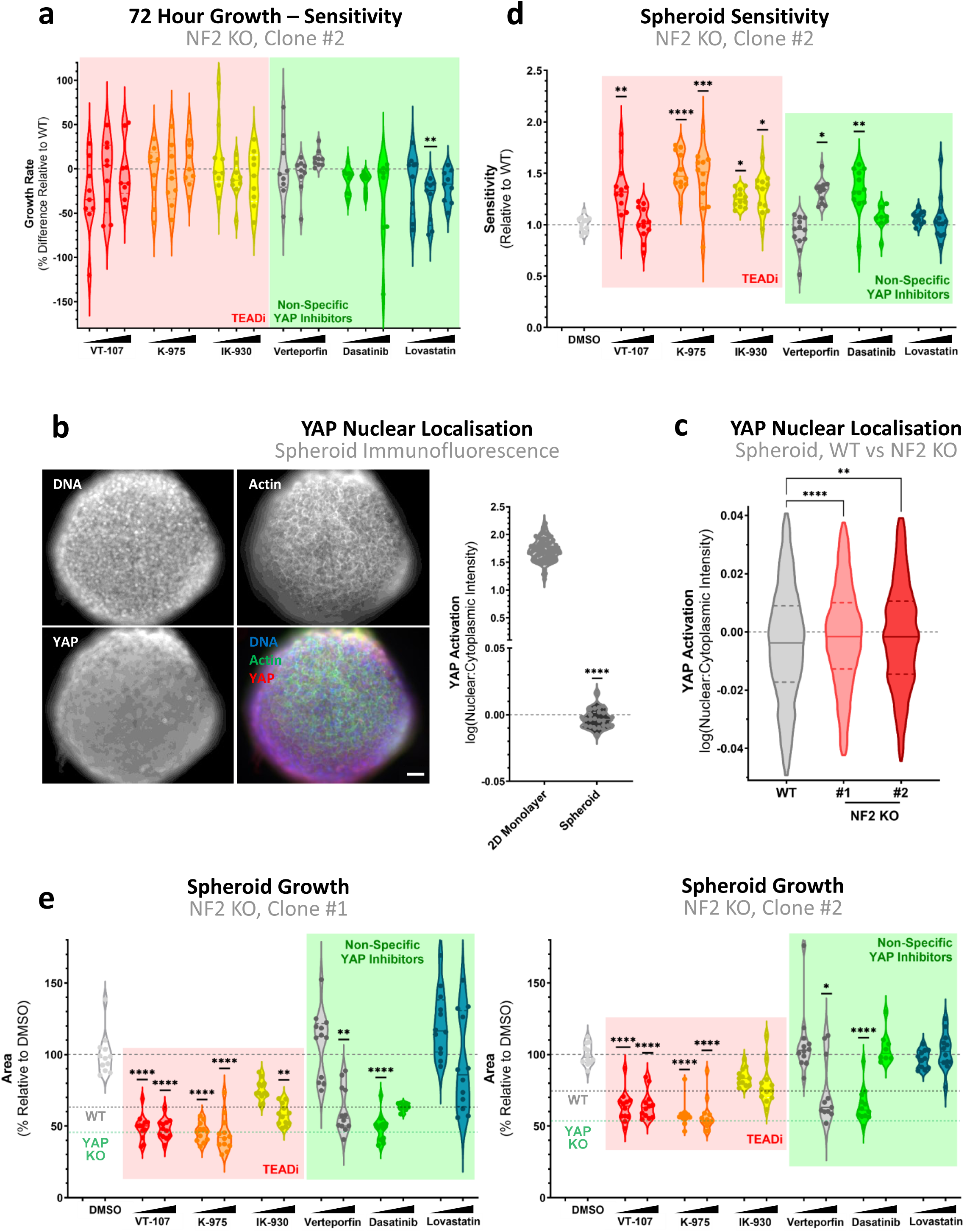
NF2 differentially modulates YAP activity in cells cultured in spheroids compared to in a 2D monolayer. **a,** Violin plot as in figure 2c representing sensitivity of NF2 KO MeT-5A cells to YAP/TAZ-TEAD inhibition, in terms of 2D proliferation. **b**, Cross-section of spheroids at day 7 (left), with labelled actin (phalloidin, green), nuclei (DAPI, blue) and YAP (red). Scale bar = 50 µm. Quantification of nuclear:cytoplasmic ratios of YAP in spheroids vs cells cultured in 2D (right) reveals a significant decrease in nuclear YAP in MeT-5A spheroids. Individual points represent median ratios across well of DMSO treated cells (2D monolayer), as in figure 1d, or median ratios within an individual spheroid (spheroid). **c**, YAP nuclear localisation analysed displayed in violin plots, from confocal images of spheroids as in (**b**) in WT (grey), NF2 KO clone #1 (light red), and clone #2 (dark red) MeT-5A cells. **d**, Sensitivity of NF2 KO spheroids (decrease in spheroid size, compared to WT spheroids) after treatment with various TEADi (red), or non-specific YAP inhibitors (green), compared to DMSO (grey). **e**, Spheroid growth of NF2 KO clones upon TEADi (red) or non-specific YAP inhibition (green) normalised to DMSO control. Dotted line indicates WT (red) and YAP KO (green) spheroid growth at steady state. Cells in (**a**) and spheroids in (**d**), and (**e**) were treated for 24 hours at 1.1, 3.3, and 10 µM and 1 and 10 µM respectively across 3 biological replicates. *P* values in (**a**), (**d**) and (**e**) were determined by Dunnet’s multiple comparison test, while those in (**b**) and (**c**) via Mann-Whitney U test. n.s. = Not significant, *P < 0.05, **P < 0.01, ***P < 0.001 and ****P < 0.0001.

**supplementary figure 3.**
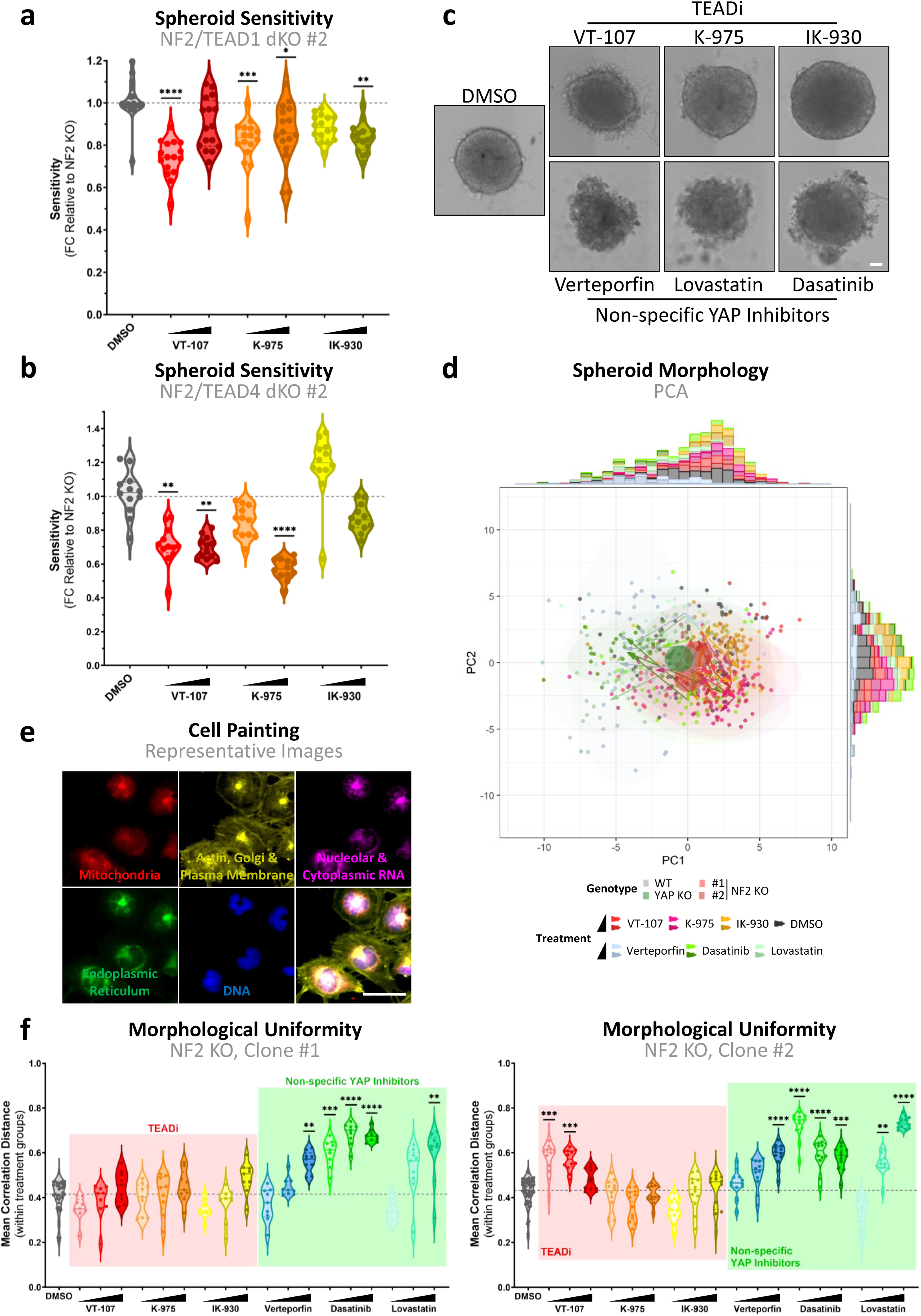
Impact of YAP-TAZ/TEAD inhibitors on spheroid morphology. **a,** Violin plot shows spheroid sensitivity to direct TEAD inhibition in NF2/TEAD1 dKO, clone #2 relative to parental NF2 KO. Points represent the decrease in spheroid growth, defined as spheroid size at day 7 relative to 24-hours, in individual NF2/TEAD1 dKO spheroids as compared to NF2 KO (n=3). **b,** Spheroid sensitivity as in (**a**) to TEAD inhibition in NF2-TEAD4 dKO, clone #2 relative to NF2 KO. **c,** Representative images of WT MeT-5A spheroids, at day 7 after treatment with 10 µM of indicated compound. Scale bar = 50 µm. **d,** PCA plot shows the distinct morphological profiles of MeT-5A spheroids after YAP/TAZ-TEAD inhibition. Spheroids were imaged and brightfield morphologies quantified, with each point representing a single spheroid. Circles represent the centre of DMSO treated spheroids, coloured according to genotype, while ellipses show the morphological spread of treatments combining genotypes. Arrows show the morphometric trajectories along increasing treatment concentrations for each genotype. **e,** Representative images of Cell Painting in WT MeT-5A cells. Indicated subcellular regions were labelled via fluorescent staining. Scale bar = 50 µm. **f,** Violin plots, as in figure 4c, showing the morphological uniformity in NF2 KO MeT-5A cells, as determined via Cell Painting, across 3 biological replicates. Each dot represents a single well. Spheroids in (**a**) and (**b**) and cells in (**d**) and (**f**) were treated for 24 hours respectively with either 1 and 10 µM or 1.1, 3.3, and 10 µM of indicated inhibitor. *P* values in (**a**), (**b**) and (**f**) were determined by Dunnet’s multiple comparison test. n.s. = Not significant, *P < 0.05, **P < 0.01, ***P < 0.001 and ****P < 0.0001.

